# PAMP: Continuous-to-Discrete Mutation Projection for Enzyme Engineering

**DOI:** 10.64898/2026.11.10.765778

**Authors:** Hengzi Wang, Zeqiu Yu, Yifei Ren, Ruizhi Yuan

## Abstract

Efficient enzyme engineering requires translating quantitative turnover predictions into specific mutations that improve catalytic activity while reducing experimental search. Here we introduce Proximal Adversarial Mutation Projection (PAMP), a gradient-guided framework that maps a differentiable turnover objective directly to single-amino-acid substitutions by combining multi-point gradient averaging with distance-penalized projection in protein embedding space. Using a substrate-conditioned Residual Predictor trained on CataPro-derived data, PAMP achieved a mean first-round predicted gain of 0.1795 log_2_ units across 2,754 enzyme–substrate pairs, with positive changes in 84.64% of cases; among the top five ranked substitutions, 99.49% of records contained at least one positive candidate. Five sequential rounds increased the mean cumulative predicted gain to 0.5631 log_2_ units, and re-scoring with an independent, non-differentiable predictor left the ordering of the methods intact while attenuating the absolute gain. PAMP provides a computationally efficient strategy for prioritizing experimentally testable enzyme mutations under small candidate budgets.

## I. Introduction

Enzymes are widely used in industrial biosynthesis [1]. For a given enzyme–substrate pair, the turnover number *k*_cat_ counts the catalytic cycles an active site completes per unit time at saturating substrate [2], [3]. Mutational scans show that most single substitutions are neutral or deleterious, with only a minority improving activity [4]. Identifying the beneficial ones experimentally requires constructing variants, expressing protein, and measuring kinetics, which is costly to repeat over a large candidate space. When throughput is limited, deciding which sites and replacements to test first becomes the central design decision, motivating computational methods that narrow the candidate set before any experiment.

Machine-learning models have increasingly provided such computational guidance. DLKcat demonstrated large-scale prediction of *k*_cat_ directly from protein sequences and sub-strate structures, establishing a practical sequence–substrate scoring framework for enzyme kinetics [2]. TurNuP extended turnover prediction to kinetically uncharacterized enzymes and reported accuracy stratified by sequence identity to the training data [5]. Subsequent models, including UniKP, CataPro, and KcatNet, further improved kinetic-parameter prediction and extended these predictors toward prospective enzyme engineering [3], [1], [6]. These studies also experimentally evaluated model-guided enzyme variants. UniKP guided the evolution of RgTAL with predicted single substitutions, CataPro combined kinetic prediction with structural and evolutionary information to engineer SsCSO, and KcatNet tested predicted *α*-glucosidase mutants [3], [1], [6].

Related design methods propose mutations in different ways. Protein language models score mutational effects zero-shot from sequence likelihood alone, without a kinetic objective [7]. kcatDiffuser instead couples graph diffusion with regressor guidance to generate mutants with higher turnover [8]. Gradient-based sequence optimization has also been applied directly. PPDE uses the gradient of a learned fitness model to build proposals inside a discrete MCMC sampler [9], and Fast SeqProp optimizes relaxed sequence parameters with straight-through gradients [10]. PAMP instead makes a fixed number of gradient evaluations around the wild-type embedding. It returns a ranked list of legal single substitutions, with no sampling, no relaxed parameters, and no generative model.

Existing *k*_cat_-guided workflows construct and score explicit candidate libraries. UniKP and KcatNet exhaustively scan single substitutions, whereas CataPro restricts positions using structural analysis and docking before enumerating replacements [3], [1], [6]. An exhaustive scan contains 19*L* mutants for an enzyme of length *L*, each requiring protein-encoder evaluation. Restricting positions reduces this cost while narrowing the sites explored.

We therefore ask whether the local response of a differentiable kinetic predictor can select a mutation directly, without building and encoding the complete library of 19*L* mutants. To address this problem, we propose *Proximal Adversarial Mutation Projection* (PAMP), which adapts gradient-based adversarial input optimization to single-substitution enzyme design [11]. PAMP uses input gradients to estimate favorable directions in the continuous protein-embedding space and subsequently projects this information onto the discrete set of legal amino-acid substitutions. The predictor defines the turnover objective, while PAMP supplies the procedure for selecting where to mutate and what replacement to introduce.

To provide a differentiable and substrate-aware optimization objective, we further develop a Residual Predictor that combines a global enzyme–substrate representation with a substrate-conditioned residue-level correction. All 19*L* legal substitution displacements are still ranked through embedding-space operations, but only the selected mutant is re-encoded for endpoint evaluation. PAMP therefore never passes the complete mutant library through the protein encoder.

## II. Methods

### A. Problem Formulation

Let *x* = (*x*_1_, …, *x*_*L*_) be a reference enzyme sequence over the 20 canonical amino acids *A*, and let *s* denote its substrate representation. Each design round returns a sequence from the single-substitution neighborhood *N*_1_(*x*) = {*x*^*′*^*∈ A*^*L*^ : *d*_*H*_ (*x*^*′*^, *x*) = 1}, where *d*_*H*_ is Hamming distance. The design objective is to maximize the predicted turnover change Δ*ŷ* (*x*^*′*^; *x, s*) over that set, in log_2_ units. PAMP does not enumerate the neighborhood, so it returns the maximizer of the projected score of Section II-C rather than of Δ*ŷ* itself. The exact one-edit constraint defines the output of each round; the substrate remains fixed. In round *r* the neighborhood is that of the current sequence *x*^(*r−*1)^, restricted to positions not yet mutated (Section II-C).

PAMP searches this neighborhood with a differentiable predictor and an adversarial selection procedure (Fig. 1). Gradients guide a continuous search and projection returns a discrete substitution. Re-encoding that substitution gives its predicted change and the input to the next round.

**Fig. 1.**
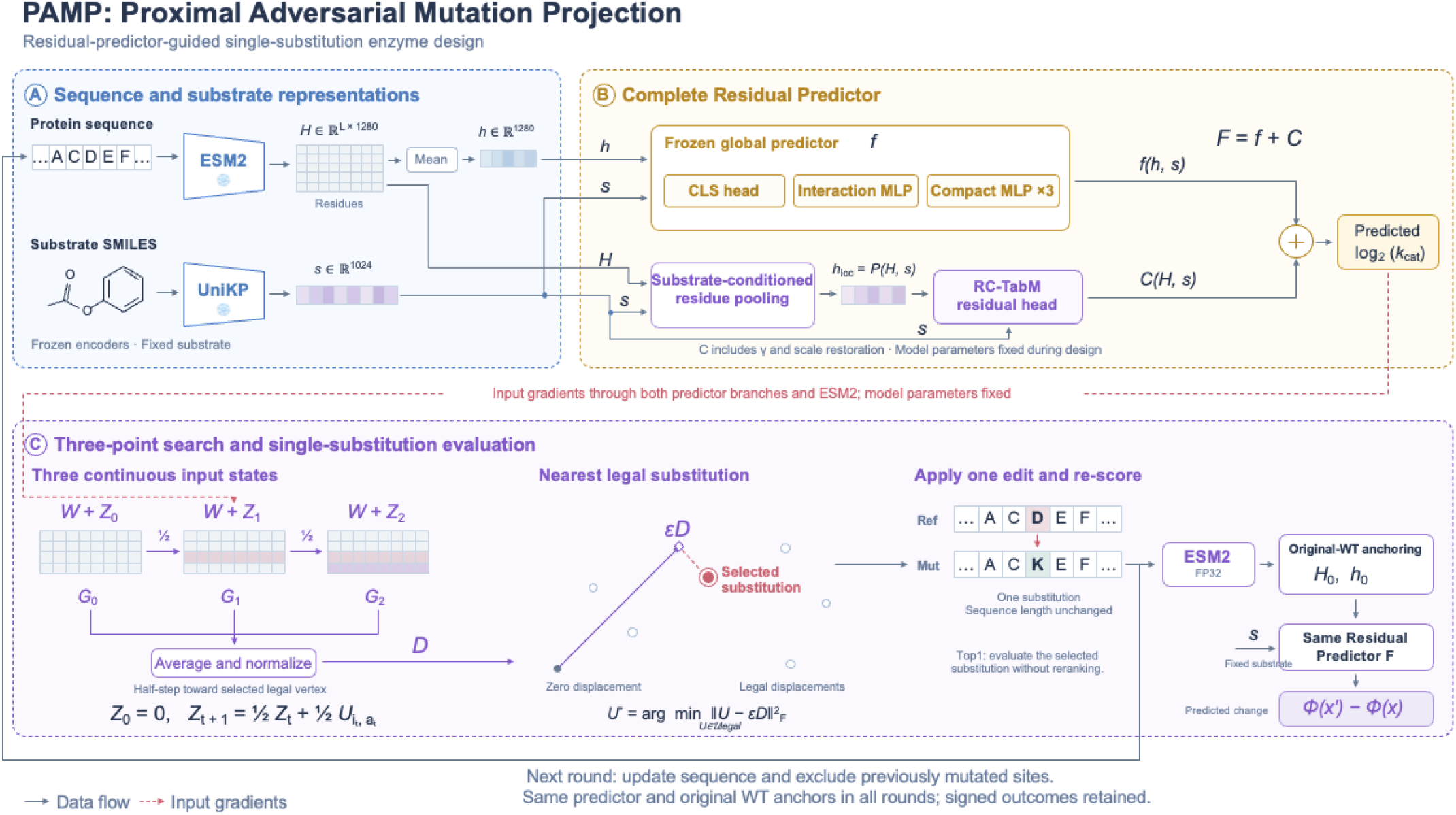
PAMP converts continuous gradient information into a single-amino-acid edit. (A) Frozen encoders provide protein and substrate representations. (B) The Residual Predictor combines a global prediction with a substrate-conditioned correction, *F* (*h, H, s*) = *f* (*h, s*) + *C*(*H, s*). (C) Continuous gradient probing averages gradients at the starting embedding and two intermediate states; discrete mutation projection maps the resulting direction to one legal substitution using the distance-penalized score; re-encoding and evaluation score the mutant with the same anchored predictor. *C* includes the learned scale *γ* and the target-scale factor *σ*_tr_.

### B. k_cat_ Residual Predictor

The complete Residual Predictor supplies the substrate-conditioned objective and input gradients for mutation design. It contains a frozen global branch, referred to as the Original Predictor, and a residue-level correction. The Original Predictor is also evaluated as a turnover-prediction baseline. We represent the enzyme using the frozen ESM2 encoder and the substrate using the UniKP SMILES Transformer [12], [3], [13]. These representations allow the objective to depend on the enzyme sequence and the substrate it acts on.

ESM2 produces 1,280-dimensional residue representations at its final layer. Their mean supplies the protein vector: for a sequence encoded in chunks *x*^(*c*)^ of lengths 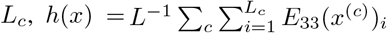 with *L* = ∑_*c*_ *L*_*c*_. The substrate encoder produces a 1,024-dimensional vector. The CLS branch projects the protein and substrate vectors to a common feature space and combines them with a learned CLS token through self-attention. A scalar head maps the resulting CLS representation to the turnover prediction. This branch learns a joint summary of the two inputs.

To complement the attention-based summary, an interaction branch explicitly combines the projected protein and substrate vectors *p* and *q*. Their concatenation, elementwise product, and absolute difference form the input *u* = [*p*; *q*; *p ⊙ q*; |*p − q* |] *∈* ℝ^2048^. An MLP maps these joint features to a turnover score. Three compact MLPs provide additional predictions from the concatenated, standardized protein and substrate vectors. The ensemble combines these branches to form the differentiable objective used by PAMP.

Let *f*_*C*_, *f*_*I*_, and *f*_*M,j*_ denote the inverse-standardized log_2_ outputs of the CLS, interaction, and compact heads. The predictor is

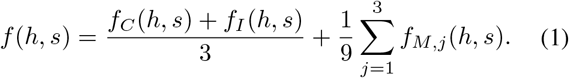

Encoder and predictor parameters remain fixed during mutation design, so optimization acts on the sequence input.

#### 1) Residue-Conditioned TabM

Substrate recognition and catalysis depend on residue interactions whose relevance varies with the substrate. Substrate-conditioned attention therefore assigns weights to residue features for each enzyme–substrate pair, producing a substrate-specific summary that complements the global protein mean.

We extend the Original Predictor with a local correction branch, termed Residue-Conditioned TabM (RC-TabM), which receives the full residue representation *H ∈* ℝ^*L×*1280^. Additive attention [14], conditioned on the substrate representation, assigns substrate-dependent weights to projected residue features while masking padding positions. The resulting pooled representation *h*_loc_ = *P* (*H, s*) is concatenated with the substrate representation and passed to a TabM head, which produces multiple residual estimates through parameter-efficient ensembling [15].

The complete Residual Predictor adds the averaged local correction to the frozen global prediction:

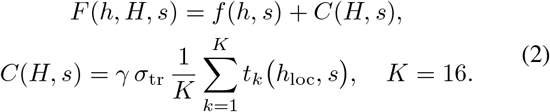

Here, *t*_*k*_ is the standardized residual from member *k, σ*_tr_ is the training-target standard deviation, and the learned scalar *γ* controls the local correction. Initializing *γ* = 0 recovers the Original Predictor; training then updates only RC-TabM and *γ*, learning substrate-dependent corrections while preserving the frozen global branch.

### C. Adversarial Mutation Search

Let *x*^(0)^ be the original enzyme of length *L*, let *x*^(*r−*1)^ be the sequence entering round *r*, and let 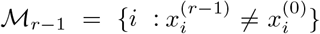 be the positions already mutated (*ℳ*_0_ = ∅). Let *W* ^(*r−*1)^ contain the ESM2 input residue embeddings of *x*^(*r−*1)^ and *V*_*a*_ the embedding of amino acid *a*. Let 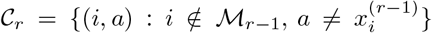 be the admissible substitutions. For (*i, a*) *∈ C*_*r*_, define the legal displacement *U*_*i,a*_ as a matrix that is zero except at residue *i*, where it equals 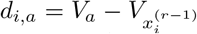. Let *H*(*W* ) and *h*(*W* ) denote the live layer-33 residue representations and their mean for input embeddings *W* .

The original wild-type anchors *h*_0_ and *H*_0_ are shared across rounds and match the training representations. The global anchor *h*_0_ comes from the mean-embedding cache; the residue anchor is computed live as *H*_0_ = *ρ*_16_(*H*(*W* ^(0)^)), where *ρ*_16_ casts to FP16 and back to FP32 to match the training residue cache. During design, precomputed caches supply only *h*_0_ and the substrate representation; intermediate states and selected mutants are encoded live. The anchored sequence score and round-*r* objective are

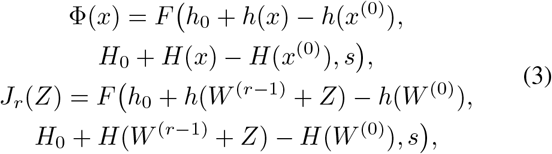

so that *J*_*r*_(0) reproduces Φ(*x*^(*r−*1)^) up to FP16 rounding. All rounds use the same Residual Predictor and the same original wild-type anchors, and the representations and gradients are recomputed around *x*^(*r−*1)^ in each round. Input gradients propagate through both the frozen global branch and the substrate-conditioned residue correction, so the search optimizes the complete Residual Predictor. We evaluate this gradient at three embedding states.

Initialize *Z*_0_ = 0 and compute *G*_0_ = *∇*_*Z*_*J*_*r*_(*Z*_0_) over residue positions. For *t* = 0, 1, choose the legal single-mutant vertex with the largest unpenalized directional score and take a half step towards that vertex:

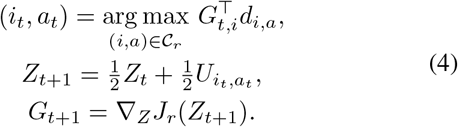

Average the starting-point gradient and the two intermediate-state gradients, then normalize over all residue positions:

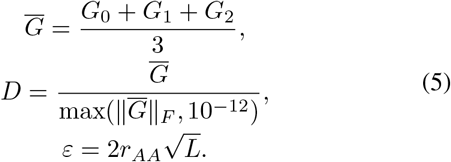

Here *r*_*AA*_ is the lower median of the pairwise input-embedding distances among the 20 canonical amino acids.

Amino-acid substitutions can alter residue interactions and conformational dynamics, making their effects on catalytic activity likely to be nonlinear and context dependent [**?**]. By averaging gradients of the substrate-conditioned predictor at the starting embedding and two adaptive intermediate states, PAMP incorporates changes in the model-predicted response along a finite perturbation path, reducing reliance on a single local linear approximation when ranking legal substitutions.

To turn the averaged direction into a substitution, PAMP balances its directional score against the squared embedding displacement:

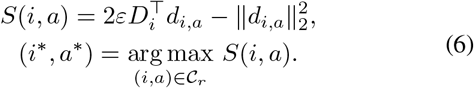

This final score is equivalent to minimizing score is equivalent to minimizing 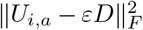, after dropping candidate-independent terms.

The distance penalty introduces a locality prior motivated by preserving the residue interactions that support enzyme structure and catalysis. Applied in ESM2 input-embedding space, it reduces the relative score of candidates with large representational displacements while retaining the substrate-conditioned directional signal. Among candidates with similar directional support, smaller displacements are preferred. This prior is imposed deterministically by the projection rule, and the matched Top1 ablation supports its practical value (Table II).

Once a substitution has been selected, the resulting sequence *x*^(*r*)^ is re-encoded to evaluate the discrete design. The predicted change of round *r* is Δ*ŷ*_*r*_ = Φ(*x*^(*r*)^) *−* Φ(*x*^(*r−*1)^), and the cumulative change is Φ(*x*^(*r*)^) *−* Φ(*x*^(0)^), which equals Δ*ŷ*_1_ in the first round. The quantity 2^Δ ŷ^ gives the predicted fold change.

For sequential design, each round selects one substitution outside ℳ _*r−*1_. The exclusion applies to both the intermediate vertex selection and the final projection, which prevents repeated positions and reversions. After *r ≤ L* rounds the sequence has exactly *r* substitutions relative to *x*^(0)^.

Every round therefore returns exactly one legal substitution. Round *r ≤ L* admits 19(*L − r* + 1) *>* 0 candidates, and this finite set has a maximizer of *S*, with ties broken by taking one of them. The guarantee concerns legality only: it does not ensure that Δ*ŷ*_*r*_ *>* 0 in any round, and it does not ensure a global optimum of the neighborhood.

## III. Experimental Setup

All mutation-search methods and ablations use the complete Residual Predictor on the same 2,754 filtered test enzyme– substrate pairs. Single-round comparisons assess proposals under one- and five-candidate budgets, ablations examine the search components, and sequential experiments examine repeated application for up to five rounds.

### A. Dataset Preparation

We trained the predictor using the CataPro-curated enzyme kinetics dataset, which draws on BRENDA and SABIO-RK [1], [16], [17]. It contains 27,658 enzyme–substrate records, 13,211 unique normalized protein sequences, and 3,302 sub-strates represented by canonical isomeric SMILES. Each record associates a sequence and substrate with a positive turnover number. Enzyme–substrate pairs were randomly assigned to training, validation, and test sets in an 80:10:10 ratio, yielding 22,126, 2,766, and 2,766 records, so a protein sequence associated with several substrates can contribute records to more than one partition. The internal test set is therefore in-distribution with respect to enzyme sequence, and every model in Table I is trained and evaluated under this same protocol. The sequence-unseen evaluation below provides the sequence-disjoint counterpart.

**Table I.** Turnover prediction and predictor ablation. RMSE is reported in log2 units. The internal test set shares enzyme sequences with the training partition; the BRENDA records contain no sequence seen during training.

| Predictor | RMSE ↓ | Pearson *r* ↑ |
|---|---|---|
| *Internal test set* (*n* = 2,766) |  |  |
| Extra Trees | 2.9754 | 0.7998 |
| CLS head | 3.2385 | 0.7621 |
| Interaction MLP | 3.2288 | 0.7746 |
| Two-head average | 3.1004 | 0.7847 |
| Original Predictor (ESMC) | 3.1848 | 0.7742 |
| Original Predictor (ESM2) | 3.0610 | 0.7891 |
| Residual Predictor (RC-TabM) | **2.9434** | **0.8075** |
| *BRENDA, unseen enzyme sequences* (*n* = 3,794) |  |  |
| Extra Trees | **4.3864** | 0.5012 |
| Residual Predictor | 4.4279 | **0.5293** |

A further evaluation assesses prediction on enzymes whose exact protein sequences are absent from the entire CataProderived source, drawn from the dataset used in DLKcat [2]. This sequence-unseen evaluation contains 3,794 enzyme– substrate records. The Residual Predictor and Extra Trees are evaluated with their trained parameters fixed; model fitting and checkpoint selection use the internal partitions. The primary mutation experiment uses 2,754 enzyme–substrate pairs from the fixed CataPro-derived test partition of 2,766 records, retaining enzymes longer than 80 residues to reduce potential outlier behavior associated with unusually short sequences. The retained cohort contains 2,391 unique enzymes ranging from 103 to 2,432 residues, with a median of 381 and an interquartile range of 299 to 498. Each pair is evaluated separately with its own substrate, and every method uses all 2,754 records.

### B. Implementation Details

Protein and substrate representations were computed using frozen pretrained encoders, with ESM2-650M [12] used for protein sequences and the pretrained UniKP substrate encoder [3] used for SMILES representations. Long proteins were encoded in nonoverlapping chunks of at most 1,022 residues and summarized by a residue-weighted mean. The encoder comparison used ESMC, averaging its final-layer residue representations into a 1,152-dimensional protein vector [18]. Turnover values were expressed in s^*−*1^, transformed to log_2_, and standardized using training-set statistics, as *y* = log_2_(*k*_cat_*/*1 s^*−*1^) and *z* = (*y − µ*_tr_)*/σ*_tr_, where *µ*_tr_ and *σ*_tr_ are the training-target mean and standard deviation. The Original Predictor’s heads were trained with AdamW [19] to minimize standardized-target mean squared error. Optimizer settings are given in the Supplementary Information alongside those for RC-TabM. Feature standardization used training data, predictions were inverse-standardized for evaluation, and the branch weights were fixed as defined in Methods.

Residue-Conditioned TabM (RC-TabM) was trained using the same data partitions and cached mean protein and substrate features, together with the complete residue-level representations extracted from frozen ESM2 [12]. RC-TabM applies substrate-conditioned additive attention [14] to the residue representations and uses a TabM head for parameter-efficient ensemble prediction [15]. The RC-TabM branch and scalar residual gate were optimized with AdamW [19] using the average member-wise squared error, while the Original Predictor remained frozen; the final checkpoint was selected according to validation RMSE. The Supplementary Information provides the detailed RC-TabM architecture, training settings, and comparisons with Mean-TabM and Standalone CondPool-TabM.

Following UniKP’s regression framework, we used Extra Trees as a prediction baseline [3], [20], concatenating the 1,280-dimensional ESM2 protein representation and the 1,024-dimensional UniKP substrate representation into 2,304 features. Extra Trees serves as a turnover-prediction baseline and, in Section IV-D, as an independent scorer for the designs; all mutation-search objectives use the complete Residual Predictor.

For mutation selection, Random provides an unguided sub-stitution baseline, whereas HotFlip ranks candidate substitutions using the gradient evaluated only at the reference sequence [11]. The remaining methods form a 2 *×* 2 design over the two components of PAMP, namely multi-point gradient averaging and the squared embedding-distance penalty. *Single-gradient* uses only the reference-point gradient while retaining the penalty. *PAMP (no penalty)* averages gradients over the three embedding states but drops the penalty. *PAMP (with penalty)* is the complete method and combines both components. HotFlip occupies the remaining cell, using the reference-point gradient without the penalty. Elsewhere in this paper, PAMP without a qualifier refers to the penalized variant. Top1 denotes the setting in which only the highest-ranked substitution is selected in each round.

### C. Evaluation Metrics

Prediction experiments use RMSE and Pearson correlation to assess numerical error and linear association with recorded turnover. RMSE is reported in log_2_ units. The primary design metrics across the *N* = 2,754 test pairs are the record-weighted mean predicted change 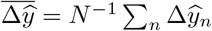 and the fraction of positive changes *p*_+_ = *N* ^*−*1^∑_*n*_**1**[Δ*ŷ*_*n*_ *>* 0]. All signed outcomes enter these summaries. The negative fraction is *p*_*−*_ = *N* ^*−*1^ ∑_*n*_ **1**[Δ*ŷ*_*n*_ *<* 0]. The reported predicted fold change is the geometric mean 2^Δ*y*^. For two-round design, the second-round increment is measured from the first-round mutant, and cumulative change is measured from the original reference.

Method comparisons are paired at the record level. Let *d*_*n*_ be the difference in predicted change between PAMP and a baseline on record *n*. We report 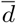 with a 95% confidence interval from a cluster bootstrap that resamples the 2,391 unique enzymes, so records sharing an enzyme are resampled together. We also report the share of records on which PAMP is higher, selects the same substitution as the baseline, or is lower, written W/T/L. A tie is exact agreement in the selected substitution, which forces *d*_*n*_ = 0. The paired difference in positive share is Δ*p*_+_, in percentage points.

## IV. Results

### A. Prediction Performance for Mutation Search

We first evaluated the complete Residual Predictor used as the differentiable scoring model for PAMP and compared it with Extra Trees, following the regression framework used in UniKP [3], [20]. On the internal test set of 2,766 enzyme– substrate pairs, the Residual Predictor achieved an RMSE of 2.9434 log_2_ units and a Pearson correlation of 0.8075, compared with 2.9754 and 0.7998 for Extra Trees, respectively (Table I).

We further evaluated both models on 3,794 BRENDA records whose enzyme sequences were absent from the CataPro-derived source. Extra Trees obtained a slightly lower RMSE (4.3864 versus 4.4279), whereas the Residual Predictor achieved a higher Pearson correlation (0.5293 versus 0.5012). The two evaluations bracket the predictor from different directions. The internal test set shares enzyme sequences with the training partition and therefore measures in-distribution accuracy, whereas the BRENDA records contain no sequence seen during training. The drop from *r* = 0.8075 to *r* = 0.5293 indicates how much of the internal accuracy depends on having encountered the enzyme during training, and the residual correlation on unseen sequences indicates that the objective retains signal outside that regime.

### B. Mutation Design

Using the complete Residual Predictor on the fixed 2,754 enzyme–substrate pairs, a single Top1 round produced a mean predicted increase of 0.1795 log_2_ units, with positive changes in 84.64% of the records (Table II). The composition of the selected substitutions supports the plausibility of the search. Conservative exchanges dominate the most frequent choices: R *→* K and E *→* D preserve side-chain charge, V *→* I and M *→* L preserve aliphatic character, and A *→* S exchanges two small residues. With K *→* Q, these six types account for 17.8% of first-round designs. Frequent charge-preserving exchanges such as R *→* K and E *→* D provide chemically interpretable examples consistent with the locality motivation of the projection; substitution matrices derived from aligned protein blocks provide an evolutionary description of amino-acid exchange preferences [21]. Tryptophan is nevertheless over-represented as a replacement and cysteine as a replaced residue, at least 16.7% and 8.6% of designs, so candidates that introduce a bulky aromatic residue or remove a cysteine would need stability screening before experimental testing. We also checked the designs structurally. Sequence recovery relative to the original reference averaged 0.9966 after one round and 0.9933 after two, matching the exact one-edit budget. Structures predicted with ESMFold [12] gave a mean TM-score [22] of 0.9872 after one round and 0.9819 after two, and mean pLDDT changed by only *−* 0.1887 and *−* 0.2872 points. The designs therefore leave the predicted fold and its confidence essentially unchanged. This is a sanity check rather than a stability assessment, because structure predictors respond weakly to single substitutions.

**TABLE II.** Ablation results for mutation search on all *n* = 2,754 enzyme–substrate pairs. Mean and Median are reported in Δ log_2_(*k*_cat_) units, and Positive denotes the proportion of records with predicted change greater than zero. Best-of-Top5 reports the best-scoring single substitution among the five highest-ranked proposals and does not represent a five-mutation sequence. Two-round Top1 results are cumulative relative to the original reference sequence.

| Method | First-round Top1 |  |  | First-round Best-of-Top5 |  |  | Two-round Top1 |  |  |
|---|---|---|---|---|---|---|---|---|---|
|  | Mean | Median | Positive | Mean | Median | Positive | Mean | Median | Positive |
| Single-gradient | +0.1098 | +0.07035 | 75.45% | +0.3607 | +0.18454 | 98.80% | +0.1628 | +0.12766 | 75.96% |
| HotFlip | +0.1089 | +0.06833 | 73.86% | +0.3672 | +0.18907 | 98.29% | +0.1471 | +0.11805 | 72.77% |
| PAMP (no penalty) | +0.1690 | +0.10154 | 83.26% | **+0.3835** | **+0.20005** | 98.98% | +0.2853 | +0.19318 | 85.00% |
| PAMP (with penalty) | **+0.1795** | **+0.10268** | **84.64**% | +0.3809 | +0.19613 | **99.49**% | **+0.3016** | **+0.19874** | **87.22**% |
| Random | −0.0847 | −0.02083 | 39.76% | +0.1403 | +0.07413 | 88.42% | −0.1712 | −0.05457 | 35.00% |

### C. Ablation Study

We conducted ablation studies on both the turnover predictor and the mutation-search procedure to examine the contribution of the main architectural and optimization components.

#### 1) Predictor Ablation

Every component of the predictor lowered RMSE on the internal test set (Table I). Averaging the CLS and interaction branches improved on either branch alone, indicating that the two pathways carry complementary information, and adding the compact MLPs to form the Original Predictor improved it further. ESM2 outperformed ESMC as the protein representation. Augmenting the frozen Original Predictor with the RC-TabM correction gave the complete Residual Predictor, lowering RMSE from 3.0610 to 2.9434 and raising Pearson correlation from 0.7891 to 0.8075, so substrate-conditioned residue-level information adds predictive signal beyond the global protein representation. Along this construction path the two largest single steps are averaging the CLS and interaction branches (*−* 0.1284 RMSE) and adding the RC-TabM correction (*−* 0.1176).

#### 2) Mutation-Search Ablation

We next examined the contributions of the main components of the mutation-search procedure on the fixed 2,754 enzyme–substrate pairs, using the complete Residual Predictor as the scoring function. The four gradient methods form a 2 *×* 2 design over multi-point averaging and the distance penalty, and Random provides an unguided substitution baseline. Under the first-round Top1 setting, HotFlip achieved a mean predicted change of 0.1089 log_2_ units (Table II), and adding the penalty alone changed this only marginally (Single-gradient, 0.1098). Multi-point gradient averaging produced the larger improvement: PAMP (no penalty) increased the mean to 0.1690, and adding the penalty on top of averaging increased it to 0.1795, with matching increases in median and positive fraction. Random substitutions instead produced a negative mean change of *−* 0.0847.

Paired tests on the same records support this ordering. At Top1, PAMP improved on HotFlip by 0.0706 log_2_ units [0.0502, 0.0914], winning on 35.6% of records, tying on 49.7%, and losing on 14.7%, with a positive-coverage gain of 10.8 points [9.0, 12.4]. Holding the penalty fixed, multi-point averaging alone accounted for 0.0697 [0.0501, 0.0896] of that difference. The penalty itself contributed 0.0105 [0.0018, 0.0203] and 1.4 coverage points [0.4, 2.3], but it left the selected substitution unchanged for 81.2% of records and won on 55.8% of the 18.8% that changed. All three intervals exclude zero while differing in magnitude by about a factor of seven, and the 2 *×* 2 design shows why: the penalty moved the Top1 mean by 0.0009 without averaging and by 0.0105 with it.

We further evaluated a small candidate-budget setting by retaining the five highest-ranked proposals from each method, evaluated independently rather than applied together. PAMP provided at least one positively scored candidate for 99.49% of the pairs, the highest coverage among the evaluated methods, with a mean best-of-Top5 change of 0.3809. The unpenalized variant obtained a slightly larger mean change of 0.3835, but that difference is not distinguishable from zero (*−* 0.0025 [*−* 0.0117, 0.0079]), and neither is the difference from HotFlip (+0.0137 [ *−* 0.0002, 0.0278]). The positive-coverage differences do exclude zero, at 1.2 points [0.8, 1.7] over HotFlip and 0.5 points [0.2, 0.8] over the unpenalized variant, as does the mean-gain comparison with Single-gradient (+0.0203 [0.0061, 0.0339]). At a five-candidate budget, the advantage of PAMP therefore lies in how many records receive at least one positively scored candidate rather than in the magnitude of the best candidate.

Together, these comparisons indicate that multi-point gradient averaging provides the dominant improvement over reference-gradient search, while the distance penalty supplies a complementary locality preference that improves Top1 candidate selection. The best-of-Top5 results further show that the ranked proposals can provide broad positive coverage when a small experimental candidate budget is available.

### D. Transfer to an Independent Predictor

The design experiments optimize and score with the same model, so we re-scored every first-round design with the Extra Trees baseline, which is not differentiable and shares no parameters or gradients with the Residual Predictor (Table III). It performs no selection: the Residual Predictor decides which substitutions are proposed and which is kept, and Extra Trees only scores that decision. Magnitudes collapse while the ordering of the methods is preserved. PAMP retains about 17% of its Top1 gain, 0.031 log_2_ units [*−* 0.003, 0.067], an interval that includes zero, and 48.0% of designs remain positive. The paired advantages survive, however, both over Random (+0.141 [0.112, 0.171]) and over HotFlip (+0.034 [0.018, 0.049]). At a five-candidate budget, where the Residual Predictor keeps the best of its own five proposals, the transferred gain reaches 0.085 [0.048, 0.122] against *−* 0.033 for five randomly drawn candidates ranked the same way. T Taken together, the preserved ranking and the five-candidate result indicate that the search is locating substitutions an independent model also rates above unguided ones, even though the absolute gain is much reduced. This bounds how far the selection depends on the guiding predictor; it is not evidence of biological validity.

**TABLE III.** Designs re-scored by an independent predictor. Residual is the guiding objective; Extra Trees shares no parameters or gradients with it and performs no selection. Values are the mean Δ log_2_(*k*_cat_) over the same *n* = 2,754 records, and Retained is the Extra Trees mean as a fraction of the Residual mean. The final row gives PAMP at a five-candidate budget

| Method | Residual | Extra Trees | Retained |
|---|---|---|---|
| PAMP (with penalty) | +0.1795 | +0.031 | 17% |
| PAMP (no penalty) | +0.1690 | +0.021 | 12% |
| Single-gradient | +0.1098 | +0.011 | 10% |
| HotFlip | +0.1089 | −0.002 | – |
| Random | −0.0847 | −0.109 | – |
| *First-round best-of-Top5* |  |  |  |
| PAMP (with penalty) | +0.3809 | +0.085 | 22% |

### E. Model Complexity Analysis

ESM2 encoding and input-gradient evaluation account for the main computational work. Per round, PAMP performs four encoder passes regardless of *L*, three gradient evaluations plus one encoding of the selected mutant, and the distance penalty adds none of them. Ranking the legal replacements then costs only *O*(19*Ld*) arithmetic for embedding width *d*, whereas an exhaustive scan requires 19*L* forward passes, which is 7,239 passes at the median cohort length of 381 residues. Matched-hardware timing against direct screening would still be needed to convert this operation count into a wall-clock speedup. The Residual Predictor contains 8,507,926 prediction-head parameters, and the projection introduces none.

Three sequential Top1 rounds over all 2,754 records took 5,346.47 s on a single NVIDIA RTX 3090 with sequences processed one at a time, giving 1,782.16 s per round and a mean of 0.65 s per record. This is a per-sequence cost rather than a batch average, and it grows with enzyme length. The figure is total elapsed time covering the three gradient evaluations, the scoring, and the encoding of the selected mutant; precomputed substrate representations and model loading are excluded, and we do not attribute time to individual operations within a round.

### F. Five-Round Sequential Mutation Design

The mean cumulative change rose at every round, from 0.1795 log_2_ units after the first to 0.3016, 0.3899, 0.4828, and 0.5631 after rounds two to five, and the share of pairs with a positive cumulative change rose from 84.64% to 88.42%. The mean per-round increment stayed positive throughout but shrank from 0.1795 to 0.0804, with a small rise in round four, and the share improving on the preceding round’s mutant fell from 84.64% to 78.98%. These results indicate that PAMP can continue to identify positively scored substitutions after multiple sequence updates, while the declining per-round gains suggest progressively diminishing improvements as the search moves away from the original sequence.

## V. Discussion

PAMP produces ranked single substitutions under fixed candidate budgets. Multi-point averaging improves candidate quality, and the distance penalty improves Top1 mean predicted gain and positive coverage. At a five-candidate budget, both variants offer similar best-candidate gains, with a coverage advantage for penalized projection. These results favor penalized projection for selecting one variant and support either variant for proposing several. Independent Extra Trees scoring preserves the method ordering, extending the comparison beyond the guiding predictor.

Existing *k*_cat_-predictor-guided workflows build an explicit candidate library and score its members, either by exhaustive single-substitution scanning or by first restricting positions with structural analysis and docking [3], [6], [1]. An exhaustive scan requires 19*L* encoder passes, whereas PAMP performs four per round regardless of *L* and ranks the legal replacements with *O*(19*Ld*) arithmetic. Against reference-gradient search [11] the averaged direction selects better candidates, and against sequence optimizers that rely on continuous relaxation or sampling, PAMP projects directly onto the legal single-substitution set, so it needs no relaxed parameters and no generative model, and every round returns exactly one legal edit.

The training data, the predictor’s error, the search approximation, and the absence of experimental measurement limit these conclusions together. Measured *k*_cat_ values are scarce relative to sequence space, and matched wild-type and mutant series are sparsely covered, so the predictor is fitted mainly to variation between enzymes rather than to the effect of a single substitution within one enzyme. That caps the precision any design method built on it can reach, and denser mutant kinetics for the same enzyme–substrate pair would raise the cap without changing the projection. The same scarcity shapes where the gains apply: the design cohort is in-distribution with respect to the training sequences, and sequence-based kinetic predictors lose accuracy as similarity to the training data decreases [5], so these gains characterize enzymes of the kind the predictor was fitted to. Most consequentially, no measured kinetics support any of the designs, so every conclusion here concerns the learned objective. The structural check shows only that the predicted fold survives, and the substitution composition reflects model preference. Candidates that introduce a bulky aromatic residue or remove a cysteine would warrant a stability filter sensitive to single substitutions before being built. We plan to measure the kinetics of top-ranked candidates as conditions allow, the step that would settle whether the predicted gains are real.

More broadly, PAMP connects differentiable protein-function objectives to explicit sequence edits through gradients with respect to the encoder’s input embeddings. An affinity predictor conditioned on a ligand or binding partner could guide optimization toward higher predicted p*K*_*D*_, while a stability predictor could guide optimization toward higher predicted melting temperature *T*_*m*_. In both cases, multi-point gradient averaging and distance-penalized projection retain the same legal single-substitution procedure under an explicit edit budget.

## VI. Conclusion

PAMP converts gradients from a differentiable enzyme-function predictor into explicit single-amino-acid substitutions. Top1 gave positive predicted changes for most pairs, a five-candidate budget covered nearly all of them, and the cumulative gain kept increasing over five sequential rounds. The resulting ranked Top1–Top5 resource spans 2,391 unique enzymes and supports continuous-to-discrete gradient projection as a practical route from differentiable protein-function prediction to mutation candidates for experimental validation.

## Data and Code Availability

The CataPro dataset and associated code are available through the original study and its public repository [1]. Code, configuration files, data partitions, design-cohort membership, and candidate-level predictions for this work are available at https://github.com/Harry1001s/PAMP.

